# Comprehensive evaluation and interpretative insights of peptide-HLA binding prediction tools using explainable artificial intelligence

**DOI:** 10.1101/2025.04.10.648169

**Authors:** Wu Guojia, Liu Xiaochuan, Wang Yuting, Yang Yang

## Abstract

Accurate prediction of human leukocyte antigen class I (HLA-I) peptide binding is pivotal for immunological research, including vaccine development and immunotherapy. However, challenges such as tool performance variability, limited interpretability, and dataset quality hinder the broader applicability of existing models. Here, we conduct a comprehensive benchmarking of 17 prediction tools using a rigorously curated dataset of over 290,000 peptides across 44 HLA-I alleles. We assess model accuracy, robustness, and interpretability, incorporating explainability methods such as SHAP and LIME to elucidate prediction mechanisms. Our analysis reveals significant performance differences, with self-attention-based models (e.g., STMHCpan and BigMHC) demonstrating superior accuracy, while CapsNet-MHC_AN, a capsule network model, offers competitive performance. Models trained on eluted ligand data outperformed those using binding affinity data, highlighting the importance of high-quality training datasets. Ensemble and multi-algorithm strategies further enhanced prediction reliability. These findings emphasize the need for continued innovation in model design, the integration of diverse datasets, and the exploration of structural predictors, providing a framework for developing more accurate, interpretable, and clinically relevant HLA-I peptide binding prediction tools.

## Introduction

Human leukocyte antigens (HLAs) are critical for immune system function, facilitating the presentation of peptide fragments to T cells. HLAs are categorized into two major classes: class I (HLA-I) and class II (HLA-II). HLA-I molecules, encoded by HLA-A, -B, and -C genes, are expressed on all nucleated cells and present peptides derived from the proteasomal degradation of intracellular proteins. These peptides, typically 8–12 amino acids in length, are loaded onto HLA-I molecules in the endoplasmic reticulum and displayed on the cell surface, where they are recognized by CD8^+^ cytotoxic T lymphocytes, triggering immune responses against infected or malignant cells (Vyas et al., 2008). Accurate identification of peptides that bind to specific HLA alleles is critical for advancing immunotherapy and vaccine development. Experimental approaches, such as biochemical binding assays and mass spectrometry (MS)-based immunopeptidomics, have been pivotal in profiling naturally presented peptides, with MS offering high throughput and broader HLA allele coverage (Hoenisch Gravel et al., 2023). The resulting data, curated in databases such as the Immune Epitope Database (IEDB), provide essential resources for understanding peptide-HLA interactions, driving the need for computational tools to predict peptide-HLA binding (Vita et al., 2019). Such tools, particularly for identifying neoantigens derived from somatic mutations in cancer, are central to personalized cancer immunotherapy, facilitating the identification of tumor-specific epitopes with therapeutic potential (Xie et al., 2023).

The complexity of peptide-HLA binding prediction has driven the adoption of advanced computational approaches, with neural networks emerging as a transformative tool. NNs generally outperform other methods, demonstrating superior predictive accuracy (Mei et al., 2020; Paul et al., 2020; Zhao and Sher, 2018). Deep neural networks (DNNs), with their multiple layers, effectively capture complex relationships in data and are well-suited for peptide-HLA binding prediction (Kriegeskorte and Golan, 2019). Attention mechanisms, particularly self-attention, have further enhanced DNNs by allowing models to focus on important sequence elements, capturing long-range dependencies critical for accurate binding prediction. Transformers, based on self-attention, have revolutionized sequence modeling, demonstrating significant promise in immunoinformatic (Vaswani, 2017). By integrating positional encoding and multi-head attention, Transformers analyze multiple aspects of peptide sequences and their compatibility with HLA binding grooves. We systematically evaluated 14 state-of-the-art peptide-HLA binding predictors, classifying them into three groups: (i) DNN-based approaches or DNN-matrix hybrids (including IEDBconsensus2.18 (Kim et al., 2012), NetMHCcons1.1 (Karosiene et al., 2012), NetMHCpan4.1 (Reynisson et al., 2020), and NetMHCstabpan1.0 (Rasmussen et al., 2016), MHCflurry2.0 (O’Donnell et al., 2020), and MixMHCpred3.0 (Tadros et al., 2024)) that combine pre-defined binding motifs with machine learning enhancements, (ii) attention-enhanced DNN models (including ACME (Hu et al., 2019), CapsNet-MHC (Kalemati et al., 2023), DeepAttentionpan (Jin et al., 2021), DeepHLApan (Wu et al., 2019), and DeepNetBim (Yang et al., 2021)) that refine feature extraction for peptides with complex binding patterns; and (iii) self-attention-enhanced DNNs (BigMHC (Albert et al., 2023), STMHCpan (Ye et al., 2023), and TransPHLA (Chu et al., 2022)) that leverage self-attention to process large, multi-dimensional datasets, excelling in diverse HLA allele predictions. Structure-based methods were excluded due to their reliance on homologous protein structures, which limit their applicability to less-characterized alleles. Our focus, therefore, remains on sequence-based approaches, which offer greater scalability and versatility for benchmarking.

Despite advancements in peptide-HLA binding prediction, previous benchmarking efforts have faced notable limitations that hinder a comprehensive understanding of the field (Mei et al., 2020; Paul et al., 2020; Zhao and Sher, 2018). Many evaluations relied on small, outdated datasets, limiting the generalizability and relevance of their findings. Moreover, recent tools frequently claim superior performance based on evaluations with limited comparisons and narrow datasets, raising concerns about potential biases in their assessment (Albert et al., 2023; Chu et al., 2022; Hu et al., 2019; Jin et al., 2021; Kalemati et al., 2023; Karosiene et al., 2012; O’Donnell et al., 2020; Rasmussen et al., 2016; Reynisson et al., 2020; Tadros et al., 2024; Wu et al., 2019; Yang et al., 2021; Ye et al., 2023). To address these gaps, we conducted an extensive benchmarking study that incorporates 14 state-of-the-art predictors and a newly compiled, large-scale validation dataset spanning 44 HLA-I alleles and over 290,000 peptides. This robust dataset provides a comprehensive platform for evaluation. Our study also integrates explainable artificial intelligence (AI) techniques, which are essential for interpreting machine learning models. By elucidating the underlying mechanisms of predictive tools, explainable AI techniques enhance our understanding of performance variations and model differences. We systematically analyze each predictor’s algorithm, usability, and interpretability, offering actionable insights and recommendations to guide the development of further prediction tools in this critical area of immunological research.

## Results

### The overall design of benchmarking HLA-I peptide binding prediction

To rigorously evaluate computational methods for predicting HLA-I peptide binding, we selected 14 recently published algorithms, representing a range of approaches in the field. Among these, MHCflurry2.0, NetMHCpan4.1 and CapsNet-MHC each included two distinct models, resulting in a total of 17 predictors. These methods were categorized into three groups based on their underlying architecture (**Supplementary Table S1**).

For the validation, we curated a robust dataset consisting of 292,784 peptide sequences, representing 44 HLA-I alleles, sourced from multiple databases following rigorous filtration. We assessed the performance of each method across various conditions, calculating a comprehensive set of evaluation metrics for the entire dataset, as well as for specific HLA variants and peptide lengths. In addition to evaluating predictive accuracy, we conducted an in-depth analysis of the computational efficiency of each tool, comparing runtime and resource consumption.

Furthermore, we performed a conservation analysis of sequence motifs associated with HLA-I ligands, aiming to identify commonalities and trends across the various peptide-binding predictions. For the most high-performing models, we generated instance-level explanations, allowing for a deeper understanding of their decision-making processes, as illustrated in **Figure 1**. This multi-faceted evaluation framework provides a holistic view of the strengths and limitations of current HLA-I peptide binding prediction methods.

**Figure 1.**
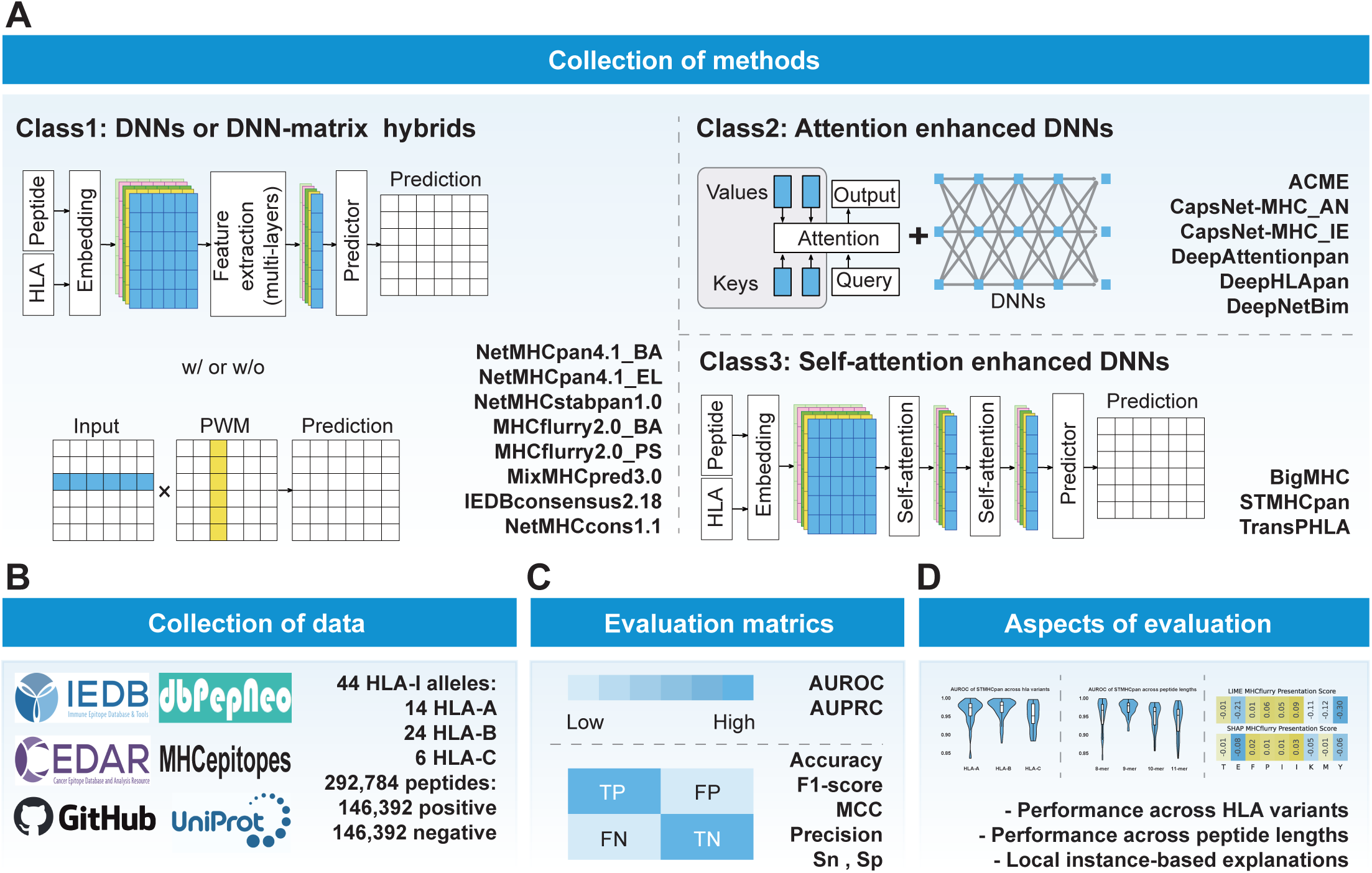
Schematic overview of the benchmarking study. **(A)** Seventeen HLA-I binding prediction methods were evaluated, including eight deep neural network (DNN)-based or DNN-matrix hybrid approaches, six attention-enhanced DNN models, and three methods incorporating self-attention mechanisms. **(B)** A diverse set of data from multiple databases was curated to assemble a comprehensive benchmark validation dataset. **(C)** The performance of the methods was assessed across a range of evaluation metrics. **(D)** The comparison also incorporated variations across different HLA-I alleles and peptide lengths, with local instance-based explanations applied to select methods for interpretability.

### Evaluation of predictor performance across the validation dataset

The validation dataset encompasses 14 HLA-A alleles, 24 HLA-B alleles, and 6 HLA-C alleles, associated with a total of 292,784 peptides of four different lengths (8-mers to 11-mers). Among these, 9-mer peptides predominate, while HLA-A and HLA-B alleles account for the majority of entries, and HLA-C alleles are relatively underrepresented (**Figure 2A, Supplementary Table S2**). Notably, all HLA-peptide pairs present in the training datasets of the evaluated predictors were excluded from our validation set. Compared with previous benchmarking studies, our dataset is notably larger and encompasses a broader diversity of HLA variants. Performance was evaluated using eight metrics: AUROC, AUPRC, Sensitivity (Sn), Specificity (Sp), Precision, F1-score, Accuracy, and MCC, calculated based on classification outcomes for the entire dataset (**Supplementary Table S3**).

**Figure 2.**
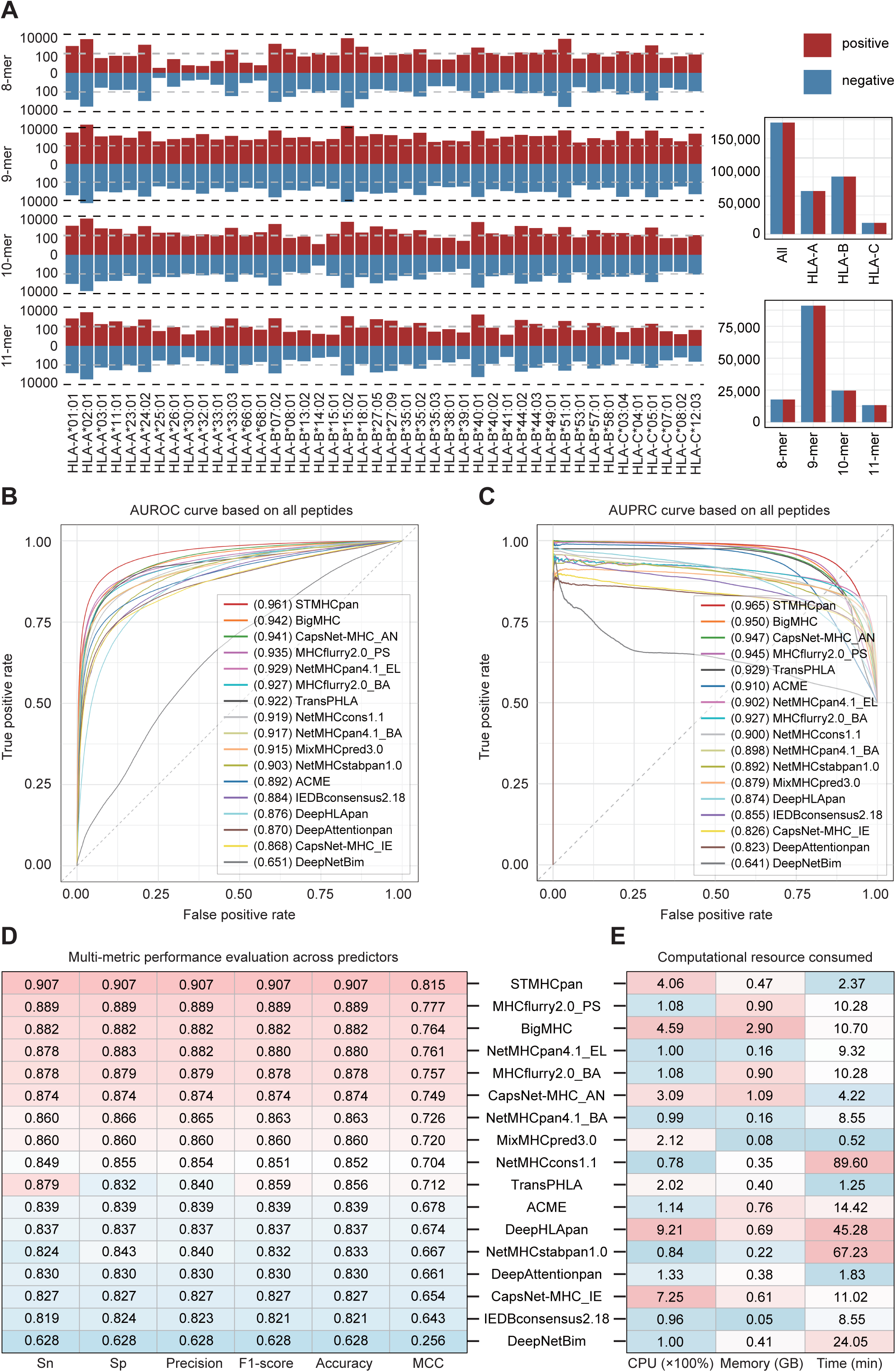
Performance evaluation of HLA-I peptide binding predictors. **(A)** Distribution of the validation dataset by HLA-I allele types and peptide lengths, represented as bar plots. **(B)** Area under the receiver operating characteristic curve (AUROC) for the performance of HLA-I binding epitope prediction methods. **(C)** Area under the precision-recall curve (AUPRC) for the performance of HLA-I binding epitope prediction methods. **(D)** Performance metrics, including sensitivity (Sn), specificity (Sp), precision, F1-score, accuracy, and Matthews’s correlation coefficient (MCC), for HLA-I binding epitope prediction methods. **(E)** Computational resource consumption analysis, showing CPU usage, physical memory consumption, and calculation time for the predictors across 4,000 selected samples.

Among the evaluated tools, STMHCpan exhibited the best performance, with an AUROC of 0.961 and an AUPRC of 0.965, surpassing other predictors across all evaluation metrics. BigMHC and MHCflurry2.0_PS also showed strong performance. While CapsNet-MHC_AN demonstrated high AUROC and AUPRC values, its binary classification outcomes were less favorable. Consistent with prior reports by the tool developers (O’Donnell et al., 2020; Reynisson et al., 2020), MHCflurry2.0_PS outperformed MHCflurry2.0_BA, and NetMHCpan4.1_EL showed superior results over NetMHCpan4.1_BA (**Figure 2B-D**). Notably, several attention-enhanced deep neural network (DNN)-based tools underperformed in our evaluation, despite claims of superior performance in their original publications (Jin et al., 2021; Wu et al., 2019; Yang et al., 2021), underperformed in our evaluation. For example, DeepNetBim yielded AUROC and AUPRC values around 0.65 and MCC values below 0.30, which were insufficient for further analysis (**Figure 2B-C**). This underperformance may be attributed to incomplete or outdated training datasets, which likely hindered the effectiveness of the attention mechanisms. This is exemplified by the performance discrepancy between CapsNet-MHC_AN, trained on Anthem’s dataset (n=539,020), and CapsNet-MHC_IE, which relied on IEDB-BD2013 (n=186,685). CapsNet-MHC_AN consistently outperformed CapsNet-MHC_IE by margins of 0.5 to 1.2 across all metrics (**Figure 2B-D, Supplementary Table S3**).

Additionally, we evaluated the computational efficiency of each method using a 4,000-entry dataset and limiting CPU usage to 10 cores. Tools such as MHCflurry2.0 and NetMHCpan4.1 demonstrated superior resource efficiency, making them suitable for use on personal computers. Conversely, tools like BigMHC and STMHCpan necessitate more advanced hardware configurations (**Figure 2E, Supplementary Table S4**).

### Performance assessment based on peptide lengths and HLA variants

We next evaluated predictor performance across peptide lengths. Irrespective of the tool or metric used, 9-mer peptides consistently achieved the highest prediction accuracy (**Figure 3A**), likely due to their prevalence as epitopes presented by HLA-I ligands and the abundance of training data for 9-mer peptides (Tadros et al., 2023). STMHCpan exhibited the highest stability, consistently outperforming other tools across all peptide lengths and evaluation metrics (**Figure 3A, Supplementary Figure S1A**). When stratifying AUROC and AUPRC scores by both HLA type and peptide length, STMHCpan achieved mean values of approximately 0.95 across all peptide lengths, outperforming the other predictors (**Supplementary Figure S1B and S1C**). Besides STMHCpan, tools such as MHCflurry2.0_BA, BigMHC, MHCflurry2.0_PS, and CapsNet-MHC_AN also demonstrated strong performance (**Supplementary Figure S1D**).

**Figure 3.**
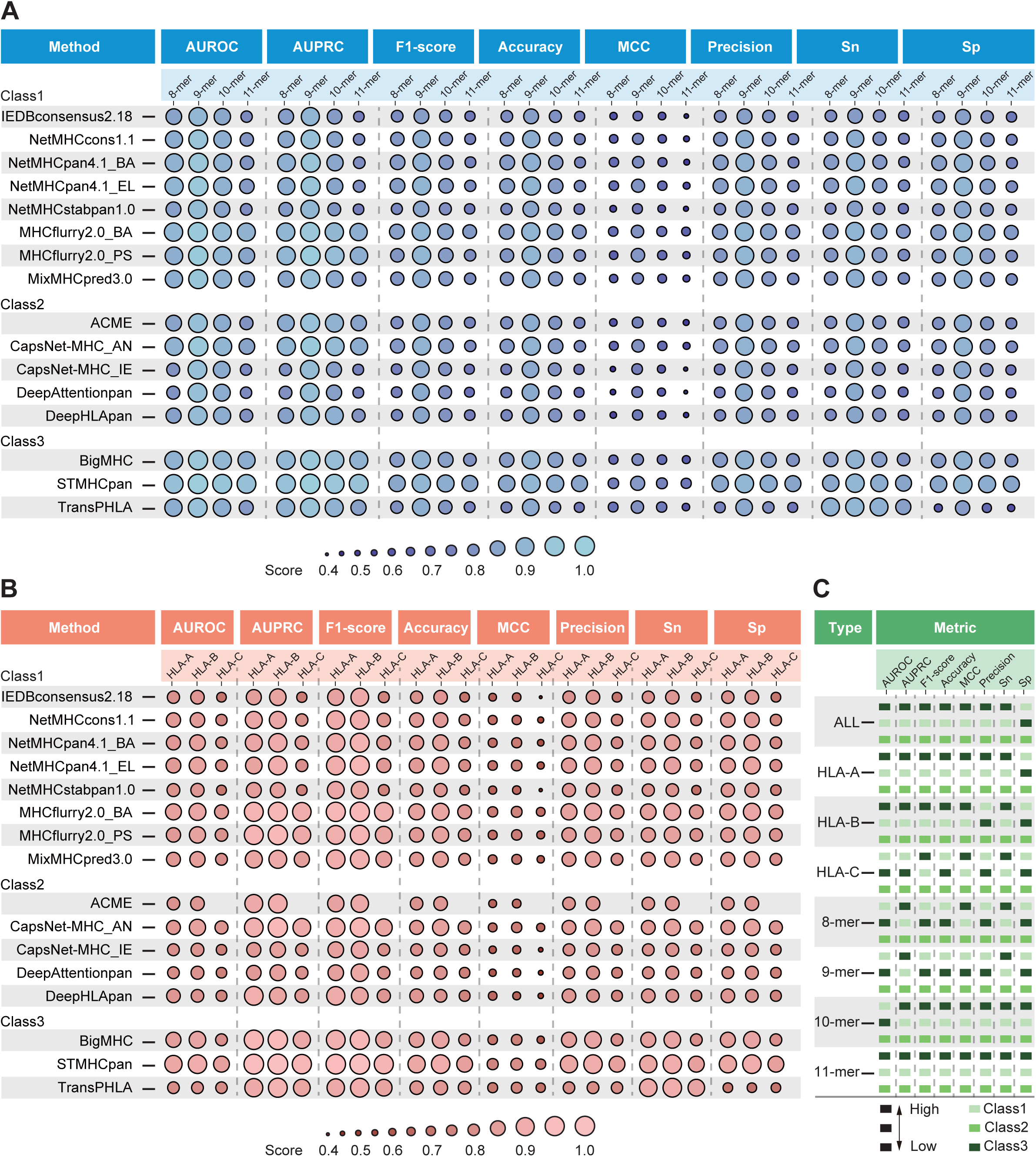
Performance of HLA-I peptide binding predictors across HLA-I types and peptide lengths. **(A-B)** Performance of the models under varying peptide lengths and HLA-I allele types, with lighter shades indicating superior performance. **(C)** Performance comparison of the three predictor classes across different scenarios, with the highest-performing class indicated in the topmost block of each group.

We then assessed performance across various HLA variants. Notably, ACME was unable to predict binding peptides for HLA-C alleles, resulting in a lack of data for this class. Overall, predictions for HLA-A and HLA-B exhibited robust prediction performance, while HLA-C alleles displayed lower performance (**Figure 3B, Supplementary Table S5**), likely due to the limited availability of antigen-binding data for this class. Certain HLA-I alleles, such as HLA-B*15:01, HLA-B*40:01, HLA-B*40:02, and HLA-B*44:03, consistently yielded strong results across multiple tools, whereas others, like HLA-C*04:01, showed significant variation in performance (**Supplementary Figure S1A, Supplementary Table S5**). STMHCpan showed superior stability and accuracy across all HLA types, with other top performers including CapsNet-MHC_AN, BigMHC, and MHCflurry2.0_PS (**Supplementary Table S5**).

We selected the top three predictors from each tool category for further analysis. From Class 1, we included NetMHCpan4.1_EL, MHCflurry2.0_BA, and MHCflurry2.0_PS; from Class 2, CapsNet-MHC_AN, ACME, and DeepHLApan; and from Class 3, STMHCpan, BigMHC, and TransPHLA. Predictors based on self-attention-enhanced DNNs outperformed those based on traditional DNN architectures. Interestingly, attention-enhanced DNN-based methods did not perform as expected, possibly due to incomplete or outdated training data. Our findings suggest that updating training datasets and adopting self-attention-based frameworks could substantially improve prediction accuracy and reliability.

### Insights into Conservation of Sequence Motifs for HLA-I ligands

Binding motifs are critical sequence patterns within peptide ligands that facilitate their interactions with the antigen-binding cleft of specific HLA-I molecules. These conserved motifs form the basis for constructing allele-specific scoring matrices, which peptide binding affinity based on characteristic positional patterns. We have compiled a dataset of 417,101 experimentally validated epitopes derived from 44 distinct HLA-I alleles. Binding motifs were generated for each HLA-I allele (**Supplementary Figure S2**).

Our analysis revealed distinct motifs for the majority of HLA-I alleles, often showing significant positional variation. However, certain alleles, such as HLA-A*26:01 8-mer (n=53) and HLA-C*07:01 11-mer (n=91), lacked sufficient peptide entries for clear motif identification. Across the majority of alleles and peptide lengths, we observed that position 2 (‘P2’), position 3 (‘P3’), and the C-terminus were consistently the most influential determinants of binding affinity. These positions align with those identified in previous studies as crucial for peptide-HLA interaction (Bouvier and Wiley, 1994; Ruppert et al., 1993). In contrast, amino acid residues at other positions showed minimal conservation across alleles.

Furthermore, we noted that binding patterns for specific HLA-I alleles exhibited clear preferences for certain amino acids, irrespective of peptide length. For example, peptides binding to both HLA-B*07:02 and HLA-C*08:02 preferentially displayed leucine (Leu) at the C-terminus (**Supplementary Figure S2A and S2B**). Additionally, closely related alleles, such as HLA-B*40:x (x=01,02) and HLA-B*44:x (x=02,03), exhibited highly similar binding motifs, suggesting shared underlying sequence preferences within these allele families (**Supplementary Figure S2H-K**). These findings contribute to a deeper understanding of the conserved sequence features that govern peptide binding to HLA-I molecules, offering insights into the mechanisms of antigen presentation and the development of allele-specific predictive models for peptide binding.

### Explainability of AI models in HLA-I peptide binding prediction

Deep learning models, while often achieving superior accuracy, are typically characterized by their lack of transparency. To address this challenge and provide greater insight into the underlying mechanisms of HLA-I peptide binding predictions, we applied explainable AI (XAI) techniques. Specifically, we focused on generating local explanations, which offer detailed insights into individual input-output pairs. These explanations are particularly valuable for understanding how models process specific instances, such as identifying critical peptide positions involved in HLA-I peptide binding predictions. In this study, we focused on local explanations for eight well-performing HLA-I peptide binding predictors across three HLA-I alleles—HLA-A*02:01, HLA-B*15:02, and HLA-C*05:01 (**Supplementary Table S6**). These alleles were selected due to their large number of 9-mer epitopes, which allowed for a robust examination while minimizing the potential impact of insufficient training data. The binding motifs for each allele are presented in **Figure 4A-C**.

**Figure 4.**
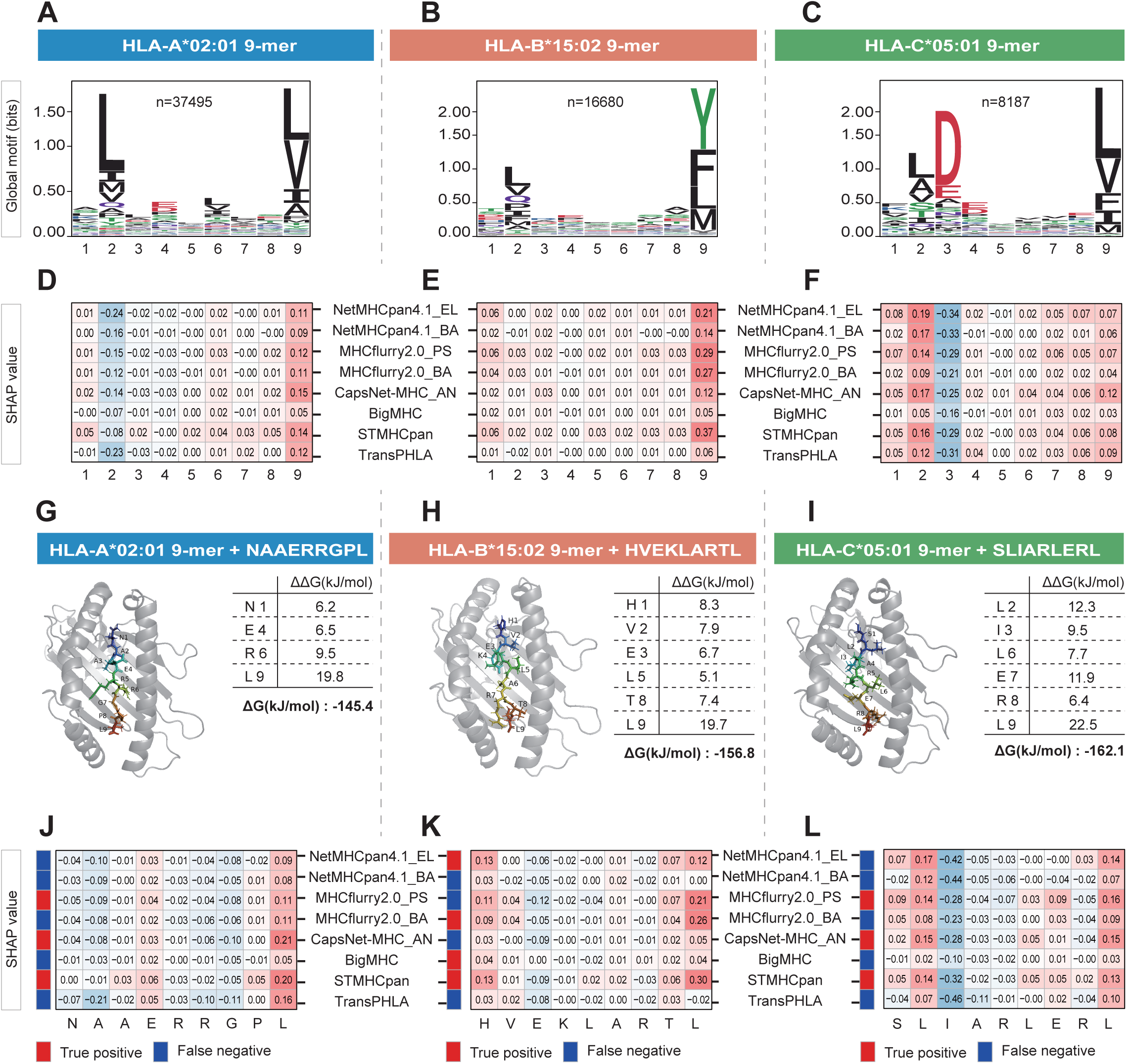
Explainable AI insights for HLA-I peptide binding predictors. **(A-C)** Binding motifs for HLA-A*02:01 9-mer* **(A)**, *HLA-B*15:02 9-mer **(B)**, and HLA-C*05:01 9-mer **(C)**. **(D-F)** Median SHAP values across 100 peptides for each allele, generated by eight different predictors. **(G-I)** peptide-HLA structures predicted by APE-Gen2.0, with ΔΔG values for each peptide residue calculated by BAlaS. Higher ΔΔG indicates greater contribution to binding, while the total ΔG of the complex reflects its stability (lower ΔG corresponds to greater stability). **(J-K)** SHAP values for each peptide residue and binary binding outcomes across the predictors.

To provide post-hoc explanations for the HLA-I peptide binding predictions, we employed two widely used explainable AI techniques: locally interpretable model-agnostic explanations (LIMIE) (Ribeiro et al., 2016) and shapely additive explanations (SHAP) techniques. For each 9-mer peptide, these methods generated attribution vectors, where each value indicated the contribution of an individual residue to the prediction. Positive attribution values represented favorable contributions to peptide binding. The median attribution values across all peptides and predictors are presented in **Figure 4D-E** and **Supplementary Figure 3A-C**. For HLA-A02:01, the patterns generated by LIME were largely consistent with those produced by SHAP. However, for HLA-B15:02, a notable discrepancy emerged at position P9, where LIME generated negative attribution scores while SHAP produced positive scores. This divergence likely reflects fundamental differences between the two XAI methods. LIME, which perturbs epitopes against a background, may overemphasize specific residues, such as tyrosine (Tyr) at P9, leading to negative scores. In contrast, SHAP incorporates a broader context of interactions, resulting in more stable and reliable attribution values. Notably, when the test dataset lacks a substantial number of Tyr residues at P9, LIME tends to yield negative patterns, while SHAP provides more balanced results. Similarly, for HLA-C*05:01, SHAP demonstrated superior consistency and reliability compared to LIME. These findings suggest that SHAP may be the more suitable technique for elucidating the predictions of HLA-I peptide binding models, offering more robust and interpretable insights, which were in consistent with prior study (Borole and Rajan, 2024).

While the predictors displayed consistent patterns across each allele for both LIME and SHAP, subtle differences were observed, likely reflecting the distinct performance characteristics of the individual models. For instance, STMHCpan, CapsNet-MHC_AN, and MHCflurry2.0_PS exhibited higher attribution scores at P9 for HLA-A*02:01, which likely contributed to their lower false negative (FN) ratios. In contrast, TransPHLA performed well at P9 but showed significant negative attribution at P2, which correlated with a higher FN ratio (**Figure 4D**). The FN ratios for these tools are provided in **Supplementary Figure 3D-F**. Notably, tools that assign higher attribution values to peptide residues generally exhibit better overall performance. However, an exception to this trend was BigMHC, which displayed a more ‘mild’ attribution profile compared to the other tools. Specifically, BigMHC did not consistently assign either extremely high or low attribution values to any residues, yet this balanced attribution did not diminish its predictive accuracy (**Figure 4D-F**).

To better understand the peptide binding predictions, we conducted a detailed analysis of three representative peptides: NAAERRGPL (bound to HLA-A*02:01), HVEKLARTL (bound to HLA-B*15:02), and SLIARLERL (bound to HLA-C*05:01). Computational methods were employed to identify key peptide residues that contribute significantly to binding, as experimental validation remains challenging. Using APE-Gen2.0, we modeled peptide-HLA (pHLA) complexes and generated multiple binding configurations for each peptide. The highest-scoring structure was selected for further analysis, employing BAlaS to compute the change in binding free energy (ΔΔG) for each peptide residue substituted with alanine. Residues exhibiting a ΔΔG ≥ 4.184 kJ/mol were considered critical for binding. This computational approach revealed binding residues that aligned with established motifs: Leu at position 9 (P9) in NAAERRGPL, Glu at position 4 (P4) in NAAERRGPL, His at position 1 (P1), Val at P2, and Thr at P8 in HVEKLARTL, and Leu at P2 in SLIARLERL. While many residues aligned with canonical binding motifs, deviations were observed, such as the presence of Glu at P3 in HVEKLARTL, which is not typically represented in the global motif, underscoring variability in binding patterns across different peptides.

**Figure 4J-K** illustrate the prediction results and attribution values derived from SHAP for each peptide, with these values generally corresponding to the computational ΔΔG. Residues with higher ΔΔG typically received higher attribution scores. The corresponding LIME results are presented in **Supplementary Figure 4A-F**. In NAAERRGPL, tools such as STMHCpan, CapsNet-MHC_AN, and MHCflurry2.0_PS assigned higher attribution values to Leu at P9 and Glu at P4, while other tools were less effective in recognizing these critical residues, resulting in discrepancies in predictions. TransPHLA, for instance, assigned a notably low score to Ala at P2, which negatively impacted the overall peptide score and led to a missed true positive outcome. The variation in tool performance was particularly pronounced in the case of HVEKLARTL, where STMHCpan assigned high attribution scores to P1 and P9, in line with their predicted contributions to binding free energy. In contrast, MHCflurry2.0_BA and NetMHCpan4.1_EL also demonstrated good performance. Notably, MHCflurry2.0_BA was more lenient in recognizing Glu at P3 compared to MHCflurry2.0_PS, leading to its correct prediction, while NetMHCpan4.1_EL outperformed NetMHCpan4.1_BA, likely benefiting from training on EL data. For SLIARLERL, MHCflurry2.0_PS showed superior performance compared to MHCflurry2.0_BA, while STMHCpan consistently produced accurate predictions.

The distribution of SHAP and LIME values for all peptides in the test dataset, along with prediction results, is publicly available at an open access data portal (**see Data Availability**). It is important to note that while the tools used in this study operate based on sequence-level content of both HLA and peptide data, the binding free energy contribution of each residue can be influenced by the peptide-HLA complex’s conformation. This conformation-dependent variability introduces a significant source of error in model predictions, as a fixed peptide-HLA combination does not always result in binding.

## Discussion

In this study, we conducted a comprehensive benchmarking analysis of 17 computational methods for predicting HLA-I peptide binding. Utilizing a rigorously curated, multi-source dataset, we ensured robust and representative testing across various conditions. Notably, we have applied model explainability techniques, such as SHAP and LIME, to offer a more detailed comparative evaluation beyond traditional metrics. The results reveal substantial variation in predictive performance, with newer self-attention-enhanced architectures showing distinct advantages. STMHCpan emerged as the top-performing tool in most settings, followed by BigMHC, while TransPHLA demonstrated high sensitivity and robust performance, highlighting the potential of these advanced architectures. Interestingly, CapsNet-MHC_AN, a capsule network-based predictor, also exhibited competitive performance, underscoring the versatility of this approach. In contrast, classical models such as MHCflurry2.0_PS and NetMHCpan4.1_EL consistently delivered good performance, reinforcing the importance of data quality and well-maintained tools. Overall, our study established a performance hierarchy, with self-attention-enhanced DNNs generally outperforming other approaches. However, the success of simpler models, particularly with comprehensive training datasets, emphasizes that continuous model refinement and careful dataset curation remain critical for high-performance predictions. This benchmarking framework, combining traditional metrics and explainability techniques, offers a standardized platform for future evaluations and advancements in AI-driven HLA-I binding peptide prediction.

Our benchmarking analysis provides significant insights into the key factors that shape the performance of HLA-I peptide binding prediction tools, revealing the complex interplay between data quality, algorithmic design, and validation methodologies. These findings not only highlight the crucial role of training data but also emphasize the need for sophisticated algorithms and robust evaluation practices in improving predictive accuracy.

The quality and quantity of training data emerged as critical determinants of model performance. Tools that incorporated eluted ligand (EL) data, derived from mass spectrometry experiments, consistently outperformed those trained primarily on binding affinity (BA) data. For instance, NetMHCpan4.1_EL, which utilizes EL data, demonstrated superior predictive accuracy over its BA-based counterpart, NetMHCpan4.1_BA, despite the use of the same underlying NNAlign_MA technique. Similarly, MHCflurry2.0, which prioritizes EL data, further underscored the value of such datasets in enhancing model performance. The success of BigMHC, which aggregated over 288,000 EL instances from various sources and incorporated a diverse set of negative examples, highlights the importance of large, high-quality datasets in boosting predictive reliability. On the other hand, tools with more limited datasets, such as CapsNet-MHC_IE and DeepNetBim, struggled to achieve similar performance, suggesting that even advanced model architectures are hindered by insufficient training data. These results underscore the need for future model development to focus on large, experimentally validated datasets, particularly those derived from EL experiments, in order to achieve more accurate and generalizable predictions.

While high-quality training data is foundational, the architecture and design of predictive models also play a significant role in determining their efficacy. Our analysis revealed that integrating multiple prediction algorithms can significantly enhance performance. For example, MHCflurry2.0, which combines outputs from antigen processing (AP) and BA predictors through a logistic regression model, outperformed each of its individual components, suggesting that hybrid approaches may better capture the complex dynamics of peptide-HLA binding. Furthermore, the potential of deep learning architectures, particularly capsule networks (CapsNet), was evident in models such as CapsNet-MHC_AN, which outperformed other advanced architectures like the transformer-based TransPHLA, despite both using the same dataset. This highlights the ability of CapsNet to efficiently model spatial hierarchies and suggests broader applicability for this architecture in HLA binding prediction. Other deep learning models, such as convolutional neural networks (CNN), recurrent neural networks (RNN), and bidirectional long short-term memory (BiLSTM) networks, could also provide valuable avenues for future research and tool development. Moreover, transformer-based models, which utilize self-attention mechanisms, showed exceptional performance even with moderate-sized datasets, suggesting that these models are particularly adept at capturing the long-range dependencies inherent in peptide-HLA binding. These findings indicate that further exploration of transformer-based architectures, in combination with self-attention mechanisms, could significantly enhance predictive accuracy in this field.

The choice of encoding methods also plays an essential role in the predictive performance of these models. Most Class 4 tools rely on one-hot encoding for peptide and HLA sequences, while others, such as those employing BLOSUM-based encodings, provide different representations that may influence feature extraction and model accuracy. Given this, the choice of encoding should be made with consideration of both the model’s requirements and its intended application. In addition, many state-of-the-art tools employ ensemble strategies, aggregating predictions from multiple models to improve overall reliability. This approach, which draws on the strengths of diverse models, enhances robustness and mitigates the risk of overfitting, demonstrating its value for increasing predictive accuracy.

## Methods

### Datasets for Benchmarking

#### (1) Assembling the Positive Set

To establish a robust external validation dataset for assessing HLA-I peptide binding prediction tools, peptide-binding data were compiled from a range of publicly accessible databases, including the Immune Epitope Database (IEDB), CEDAR, dbPepNeo2.0, and MHCepitopes (Kawakita et al., 2024; Lu et al., 2022; Vita et al., 2019). Additionally, training datasets for relevant predictors were sourced from repositories, where available. A stringent filtration process was implemented to ensure both dataset quality and independence: (i) Only peptides with experimentally confirmed binding, supported by reference sources, were retained. (ii) Peptides were restricted to naturally occurring amino acids and lengths of 8–11 residues. (iii) Duplicate peptides associated with the same HLA allele were excluded to minimize redundancy. (iv) Peptide-HLA combinations present in the training datasets of any evaluated predictor were removed to maintain the dataset’s independence. (v) HLA alleles with fewer than 250 unique peptide entries following filtering were excluded to ensure statistical robustness. The final dataset comprises 146,392 non-redundant peptide ligands across 44 distinct HLA alleles, providing a comprehensive and unbiased resource for benchmarking peptide binding prediction tools. Summary statistics are provided in Supplementary Data 1, with the full peptide list available in Supplementary Data 2.

#### (2) Assembling the Negative Set

Negative datasets were constructed by generating artificial non-binding peptides, a common practice in predictor development and benchmarking studies. Source protein sequences corresponding to the peptides were retrieved from the UniProt database. These antigen-protein sequences were segmented into peptides of lengths 8, 9, 10, or 11 amino acids, which were subsequently pooled by length. Peptides already present in the independent positive set or the training set of the evaluated predictors were excluded to maintain dataset independence. Negative peptides were then randomly selected to match the number of positive peptides for each peptide length and HLA-I allele. Although this approach may result in a small proportion of misclassified peptides—whose binding properties were not experimentally validated—the specificity of HLA-I molecules, which bind only a limited subset (∼1%) of all possible peptides from a given protein, renders the potential false-negative rate negligible relative to the overall dataset size. For example, for a 100-residue protein, one would expect approximately one true binder and around 99 non-binders, making the false-negative rate orders of magnitude smaller than the true negative count.

### Predictive performance evaluation

To assess the predictive performance of the 17 evaluated tools, we utilized a variety of standard metrics, drawn from previous studies, as all tools produce quantitative outputs, such as binding affinity (BA) scores and/or BA percentile ranks. The establishment of a standard threshold is essential for classifying antigen peptides into epitopes and non-epitopes. While some tools, such as NetMHCpan4.1 and MHCflurry2.0, provide official threshold recommendations, many others do not. To determine a threshold for each tool, we computed the median predicted score for each combination of allele and peptide length category. This approach ensures fairness and accuracy by maintaining an equal number of positive and negative samples within each category, facilitating consistent binary classification.

The metrics employed to evaluate performance included the Matthews correlation coefficient (MCC), accuracy, F1-score, sensitivity (Sn), precision, and specificity (Sp). Each metric offers a distinct perspective on model performance:

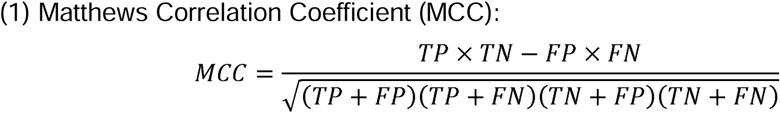

where TP, TN, FP, and FN represent the counts of true positives, true negatives, false positives, and false negatives, respectively. The MCC is a balanced measure of performance, particularly useful when dealing with imbalanced classes.

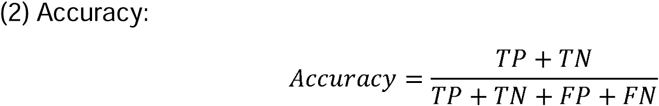

Accuracy reflects the overall performance of the model, considering both true positives and true negatives relative to the total number of cases.

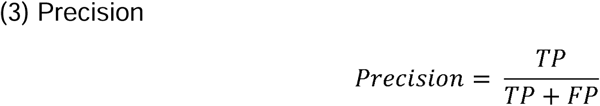

Precision measures the proportion of predicted positives that are correctly identified as actual positives.

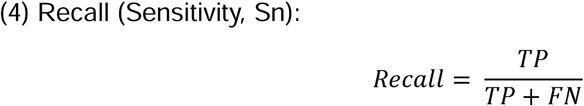

Recall, or sensitivity, evaluates the model’s ability to correctly identify true positives.

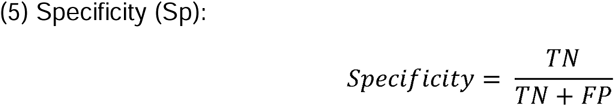

Specificity, also known as the true negative rate, measures the model’s ability to correctly identify negative instances. This metric is particularly important in contexts where the cost of false positives is high.

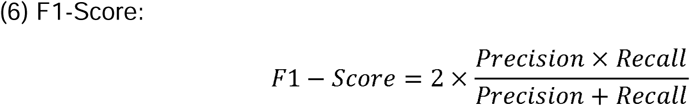

The F1-score is the harmonic mean of precision and recall, offering a balanced measure of performance, particularly in situations with imbalanced class distributions.

Additionally, we incorporated the area under the Receiver Operating Characteristic (ROC) curve (AUC) and the area under the Precision-Recall curve (AUPRC) as key performance metrics. The AUC quantifies the model’s ability to discriminate between binders and non-binders, with a value of 0.5 representing random predictions and a value of 1.0 denoting perfect predictions. The AUPRC offers a complementary measure of model performance, particularly valuable in imbalanced datasets where the precision of positive predictions is critical.

To assess the statistical significance of differences in AUC, we applied bootstrapping with replacement and used a paired t-test to compare the AUC values between predictions, determining whether the AUC of one model was significantly superior to that of another. These metrics, when considered together, provide a comprehensive evaluation of each tool’s predictive performance across varying peptide lengths and HLA alleles.

### Computational resource consumption

All experiments were conducted on a CentOS Linux 8 workstation equipped with four Intel® Xeon® Gold 6238 processors, each operating at a base frequency of 2.10 GHz, with a maximum frequency of 3.7 GHz. The system is configured with a total of 176 logical processors, organized into 22 cores per socket with hyper-threading enabled, resulting in 88 physical cores across four sockets. Each core is equipped with 32 KB of L1 data cache (L1d), 32 KB of L1 instruction cache (L1i), 1024 KB of L2 cache, and a shared 30,976 KB of L3 cache per socket. The workstation is equipped with 1.0 TiB of RAM and an additional 4.0 GiB of swap space. To ensure consistent computational performance and equitable resource distribution, each predictor was limited to utilizing a maximum of 10 CPU cores, facilitating fair allocation of system resources and reproducibility of results across all experiments.

### Instance-based explanations for HLA-I binding predictions

We utilized the MHCXAI framework to generate instance-based explanations for predictions made by HLA-I binding predictors, leveraging SHAP and LIME for their capacity to provide interpretable insights across diverse model architectures. SHAP, based on cooperative game theory, quantifies the contribution of individual peptide residues to model predictions by calculating Shapley values through iterative perturbations. LIME, on the other hand, employs local surrogate models to approximate complex predictions by assessing the effects of perturbed peptide sequences.

To address the specific input requirements of the selected predictors, we extended the MHCXAI framework to process peptide-MHC allele pairs and generate comprehensive explanations. These enhancements improved the framework’s compatibility and computational efficiency, particularly when applied to large datasets. The explanations generated by SHAP and LIME identified critical peptide residues involved in MHC binding predictions, providing valuable insights into model behavior and enhancing interpretability. This systematic approach aligns explanation methods with predictor-specific outputs while maintaining scalability and efficiency.

Eight high-performing methods—NetMHCpan4.1_BA, NetMHCpan4.1_EL, MHCflurry2.0_BA, MHCflurry2.0_PS, CapsNet-MHC_AN, BigMHC, STMHCpan, and TransPHLA—were selected for generating explanations. Given the computational resources required to apply MHCXAI methods across all peptides, a representative test dataset was carefully constructed. The dataset preparation involved the following steps:

1. Positive binding peptides for HLA-A*02:01, HLA-B*15:02, and HLA-C*05:01 were extracted from the validation dataset.
2. Peptides for which all eight methods produced identical binary predictions were excluded.
3. For each selected allele, a global position weight matrix (PWM) was derived from Supplementary Data 9, alongside the false negative (FN) ratio rank across the eight predictors.
4. Multiple random samples of 100 peptides were selected for each allele, and their PWMs and FN ratio ranks were calculated. The sets most similar to the initial PWM and FN ratio rank were selected as the final test dataset.

The detailed peptide list is provided in Supplementary Data 4.

### Validation for explanations

To evaluate the reliability of the explanations generated by SHAP and LIME, we compared the predicted important peptide residues with computational ground truth derived from changes in free energy (ΔΔG) upon mutation of individual peptide residues in peptide-MHC binding. Experimentally, alanine-scanning mutagenesis provides a precise assessment of the energetic contribution of each peptide residue to binding affinity. However, due to the resource-intensive nature of this approach, we utilized computational alanine-scanning mutagenesis via the BAlaS tool to estimate ΔΔG values for peptide residues within the binding pocket of the MHC-I complex. The pipeline began with structural predictions of peptide-MHC-I complexes generated using APE-Gen2.0, where the most stable binding conformations were selected from the predicted PDB models. Conformation stability was determined based on the quality of peptide-MHC interactions, ensuring that only the most reliable structures were included for subsequent analysis. These selected structures were then input into BAlaS, which calculated ΔΔG values for each peptide residue. Residues with ΔΔG ≥ 4.184 kJ/mol were classified as critical for binding, indicating that their mutation reduced binding affinity significantly. Conversely, residues with ΔΔG ≤ −4.184 kJ/mol were identified as enhancing binding affinity relative to the native configuration, while those with intermediate ΔΔG values were deemed to have a neutral impact. By comparing the residues identified as critical by SHAP and LIME with the computational ΔΔG results, we assessed the accuracy and biological relevance of the AI-generated explanations.

## Data availability

The binding motifs for each HLA-I allele, along with the SHAP and LIME values for all peptides in the test dataset and their corresponding prediction results, are publicly available via an open access data portal (http://bioailab.com/MHCbenchmarking).

## Code availability

The code used for this study can be obtained from Github repository (https://github.com/YY-TMU/MHCI_benchmark)

## Acknowledgements

The work was supported by the following grants: National Natural Science Foundation of China (32470721, 32100534), and Talent Excellence Program from Tianjin Medical University. We gratefully acknowledge the technical support provided by the High-performance Computing Platform of Tianjin Medial University.

## Author contributions

Guojia Wu: Writing–original draft, Visualization, Methodology, Investigation, Formal analysis, Data curation. Xiaochuan Liu: Methodology, Investigation, Formal analysis. Yuting Wang: Supervision, Resources, Project administration, Methodology, Investigation. Yang Yang: Writing–review & editing, Visualization, Supervision, Resources, Project administration, Methodology, Investigation, Funding acquisition, Data curation, Conceptualization.

---

**Supplementary Figure S1.**
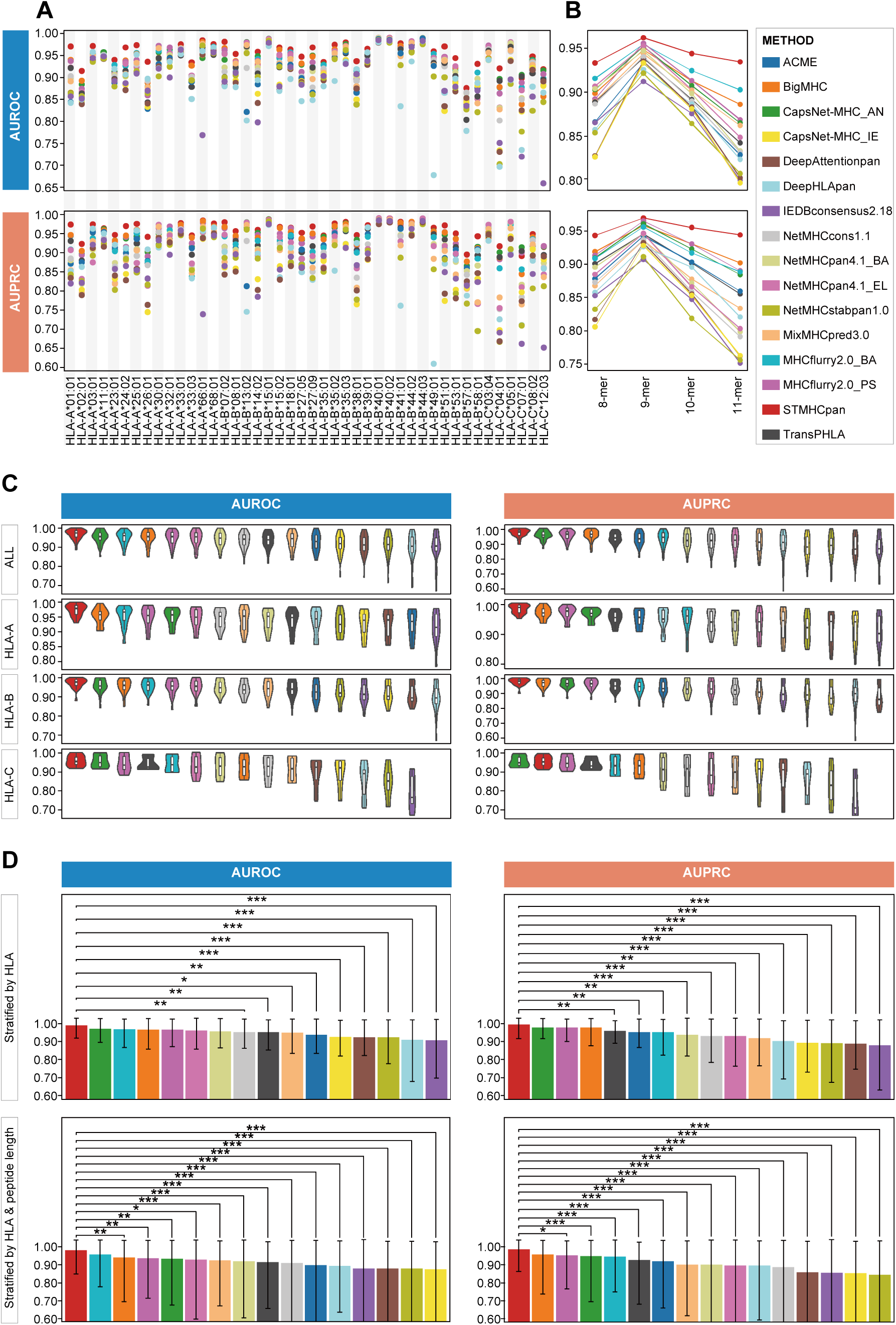
AUROC and AUPRC Performance of HLA-I peptide binding predictors. **(A-B)** AUROC and AUPRC values for each HLA-I allele and peptide length. **(C)** Violin plots overlaid with box-and-whisker plots depicting AUROC and AUPRC, stratified by allele and organized by HLA-I locus. **(D)** Mean AUROC, AUPRC, and 95% confidence intervals, stratified by HLA-I allele (n = 44) and both HLA-I and epitope length (n = 176), with adjusted two-tailed Wilcoxon signed-rank test P-values for inter-method comparisons.

**Supplementary Figure S2.**
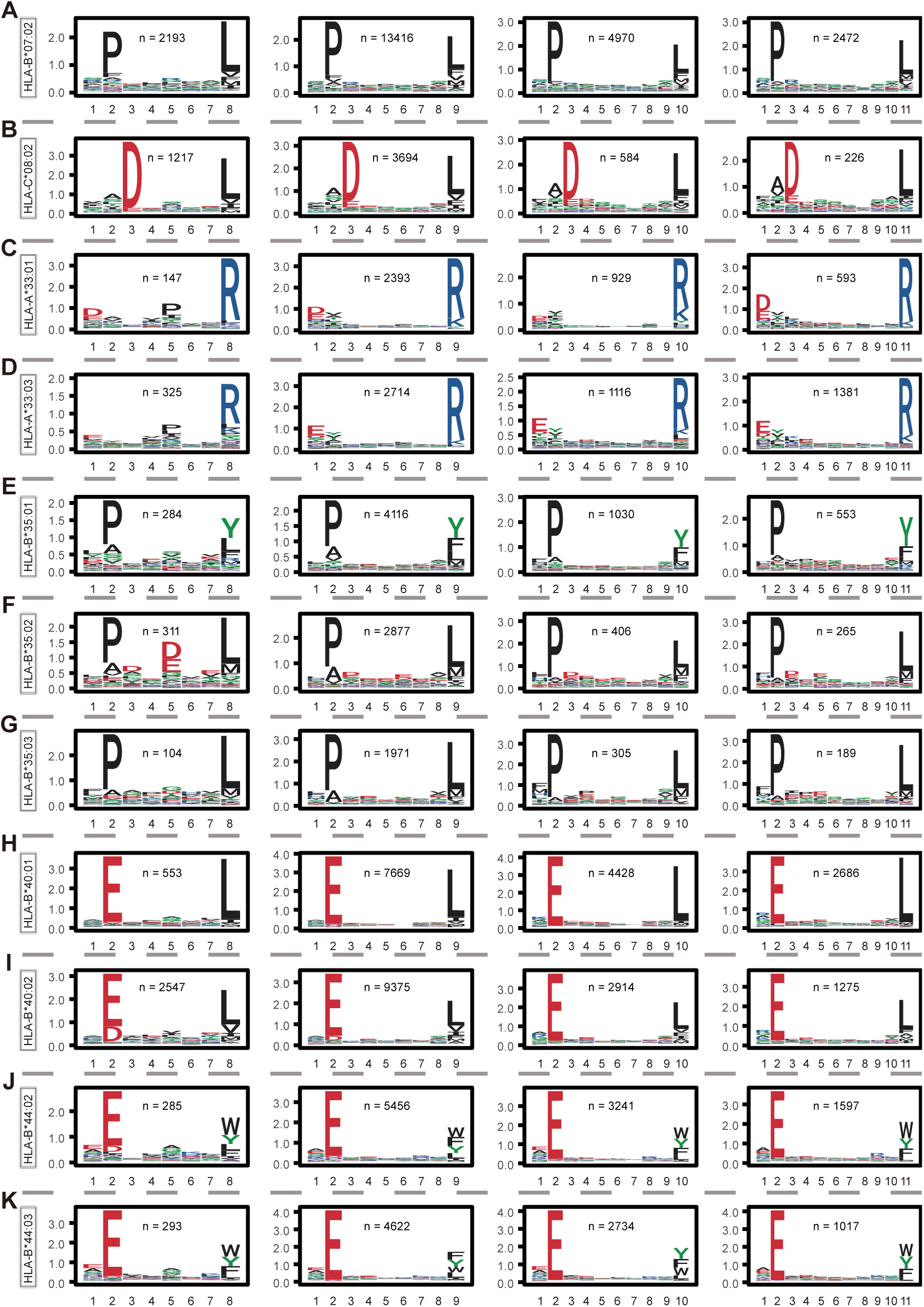
Position and residue specificity of 11 HLA-I alleles. Position and residue specificity for 8-, 9-, 10-, and 11-mer peptides of the following HLA-I alleles: **(A)** HLA-B07:02, **(B)** HLA-C08:02, **(C)** HLA-A33:01, **(D)** HLA-A33:03, **(E)** HLA-B35:01, **(F)** HLA-B35:02, **(G)** HLA-B35:03, **(H)** HLA-B40:01, **(I)** HLA-B40:02, **(J)** HLA-B44:02, and **(K)** HLA-B*44:03.

**Supplementary Figure S3.**
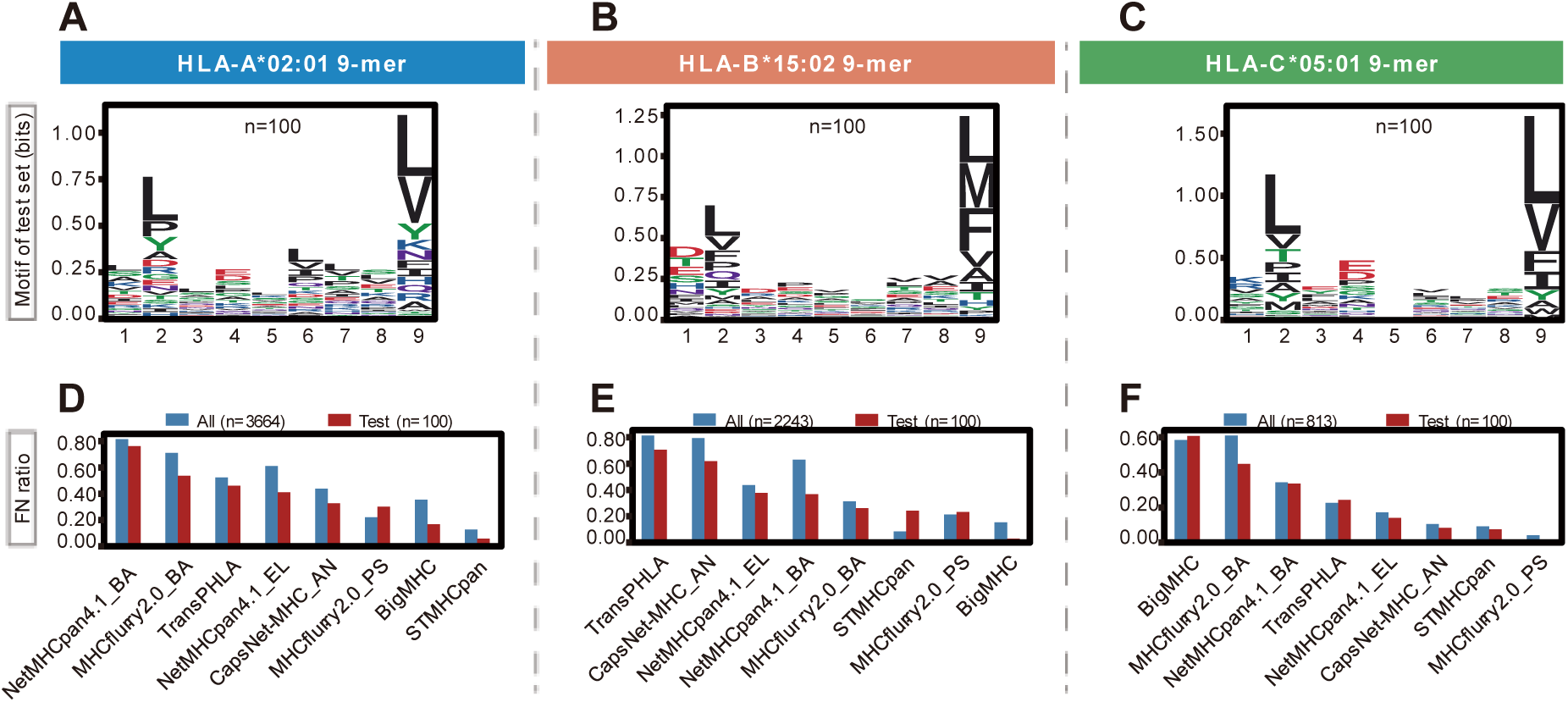
Validity of test dataset for explainable AI. **(A-C)** Binding motifs for each selected HLA-I allele in the test dataset. **(D-F)** False negative (FN) ratio of each predictor in the test dataset compared to the original dataset. A lower FN ratio indicates better prediction performance.

**Supplementary Figure S4.**
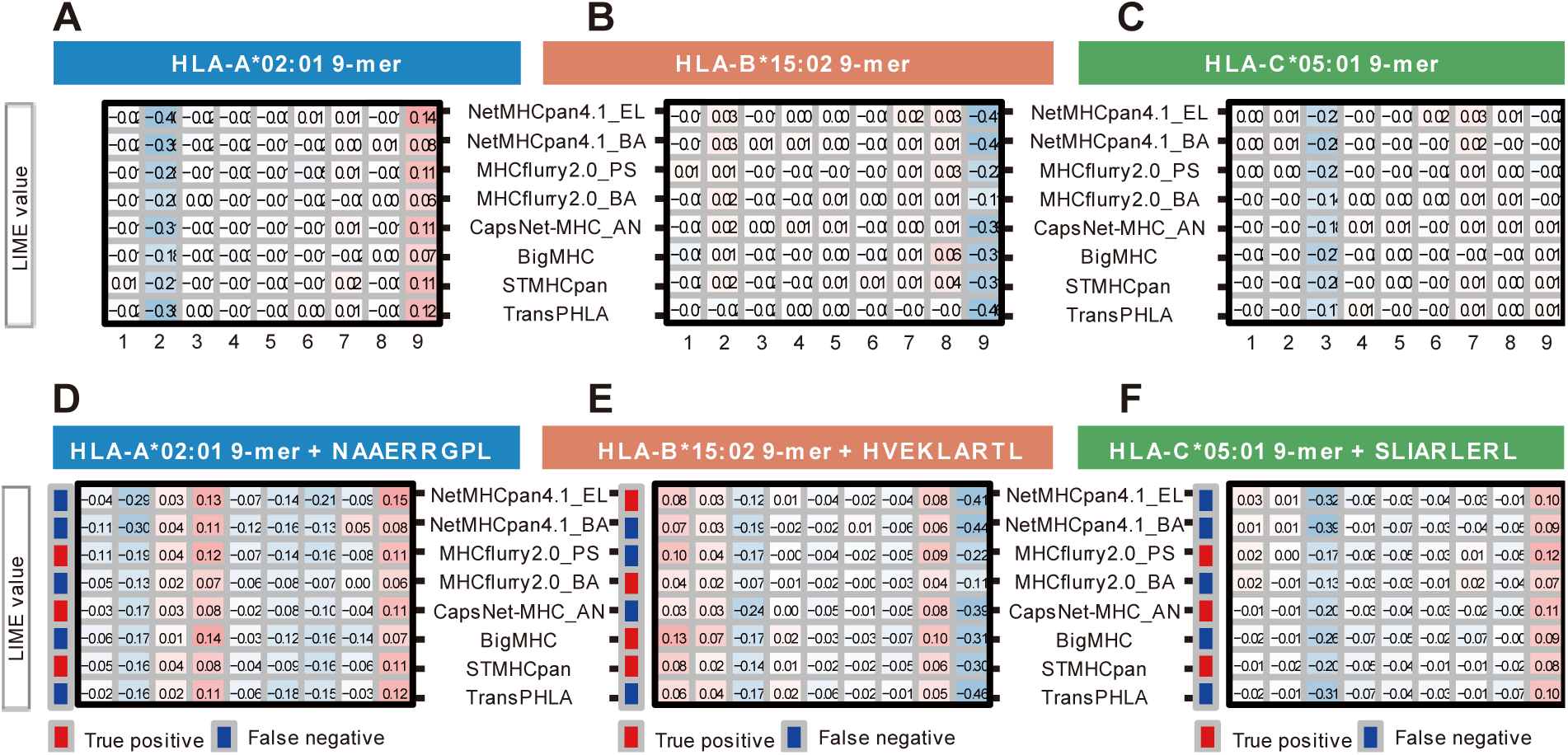
LIME values of HLA-I peptide binding predictors. **(A-C)** Median LIME values generated by eight predictors across 100 peptides for each allele. **(D-F)** LIME values for each residue of the ligand peptide and corresponding binary binding results across predictors.

